# Hydrothermal Synthesis of Clove Bud-Derived Multifunctional Carbon Dots Passivated with PVP- Antioxidant, Catalysis, and Cellular Imaging Applications

**DOI:** 10.1101/2021.11.29.470381

**Authors:** Anurag Kumar Pandey, Kamakshi Bankoti, Tapan Kumar Nath, Santanu Dhara

## Abstract

The present study reports a one-step synthesis of PVP-passivated clove buds derived carbon dots (PPCCDs) using clove buds and polyvinylpyrrolidone (PVP) as the starting precursors via the hydrothermal route. The adopted technique is facile and environmentally friendly for the production of carbon dots (CDs) with in situ PVP passivation. The study evidenced the significant modulation in optical properties of passivated CDs as compared to non-passivated ones. Structural and morphological studies evidenced the spherical PPCCDs with a diameter of ∼ 2 nm and crystalline in nature with an interlayer spacing of 0.33 nm. The PPCCDs showed excellent antioxidant activity against DPPH and superoxide anion radicals and also showed good catalytic activity for the degradation of Rhodamine-B (Rh-B) dye under the studied conditions. Their bioimaging potential was evidenced through live-cell fluorescent imaging with 3T3 and L929 cell lines.

## 1. INTRODUCTION

During the past two decades, various carbon nanostructures with different physicochemical properties have been inspected due to their distinct shapes and sizes, along with several known allotropes and structures [1]. Notably, several groups are fascinated by the astonishing properties of carbon dots (CDs) that was first reported in the process of refinement of single-walled carbon nanotubes (SWNTs) [2]. Their superior properties like photoluminescence, electron donor/acceptor ability, and photo-bleaching in comparison to traditional quantum dots or metallic nanocrystals made them the most promising in the fields of bio-labeling, free radical scavenging, catalysis, solar cells, metal ion sensors, and many more [3–9].

The selection of synthesis techniques and precursors for CDs are set a benchmark towards its applicability in various fields. There are several techniques for carbonization and fabrication of CDs such as laser ablation, combustion, pyrolysis, hydrothermal, and microwave irradiation, etc. [10–16]. Herein, the hydrothermal route is the most efficient due to its simplicity, low process cost, and controllable reaction conditions [3]. The carbon sources or precursors for CDs synthesis can be classified as either synthetic or natural in origin [5]. Among them, nature-origin carbon sources are investigated more fascinatingly because of their low-cost, nontoxic behavior, and convenience in synthesis. The nature of precursors is a crucial factor in deciding the photophysical and photochemical properties of CDs for their applicability. The quest for a novel carbon precursor at low cost, high quantum yield, and having multifunctional potential is still desirable.

Multifunctionality aspects of CDs can be attained using two adapted techniques, viz. doping and surface passivation. These techniques involve the inclusion of dopant atoms in the core and the attachment of functional groups over the surface of CDs intrinsically or extrinsically. Intrinsic passivation depends totally on carbon precursors, while extrinsic passivation uses external agents to modify the surface of CDs. Extrinsic passivation is beneficial in terms of adding new functionality in nanomaterials. It is used for the improvement of aqueous stability, surface properties, and optoelectronic properties of CDs with extensions in their application ranges. In recent years, various polymers such as PAA, PEI, PEG, and TTDDA have been used with natural or synthetic sources as passivating agents for CDs synthesis [12–16]. But the multifunctionality aspect of CDs still needs to be surveyed to verify their potential as smart nanomaterials without compromising their simplicity and toxicity. However, to our best knowledge, there is no significant study on the synthesis of polyvinylpyrrolidone (PVP) passivated CDs derived from clove buds along with their diverse applications.

By keeping sight of the aforementioned issues, one-pot hydrothermally synthesis of PVP-passivated CDs (PPCCDs) derived from clove buds is carried out. This strategy is simple, time-saving, eco-friendly, and employs an aqueous solvent in comparison to other passivation techniques reported so far [13,14]. Clove buds (*Syzygium aromaticum*) belong to the family of Myrtaceae, which is a tropical plant that grows mainly in the southern part of India. Aqueous extract of clove buds is used for the treatment of dyspepsia and gastric irritation and is also reported as a very good antibacterial and antioxidant agent [17,18]. PVP is a non-toxic, water-soluble, biocompatible polymer used as a surface stabilizer in the synthesis of nanomaterials, pharmaceuticals, the food industry, and biomedicine [19,20]. So, due to PVP passivation with nitrogen as a dopant, these CDs can enhance their multifunctionality aspects with diversity in applications. In this article, the physicochemical and optoelectronic properties of synthesized PPCCDs were analyzed. Furthermore, our study also inspects the multifunctional facets of PPCCDs in various applications such as free radical scavenging over DPPH and superoxide anion assay, catalysis over the degradation of organic dye Rh-B in presence of sodium borohydride (NaBH_4_) as a reducing agent, and bioimaging using the 3T3 (mouse embryonic fibroblasts) and L929 (mouse fibroblast) cell line.

## 2. MATERIALS AND METHODS

The floral clove buds were purchased from a local grocery store of the Indian Institute of Technology Kharagpur, West Bengal, India. Dimethyl sulfoxide (DMSO), 3-(4,5-dimethylthiazol-2-yl)-2,5-diphenyltetrazolium Bromide (98%) (MTT), Butylated hydroxytoluene (BHT), Nitro blue tetrazolium chloride (NBT), and 2, 2-diphenyl-1-picrylhydrazyl (DPPH) were procured from Sigma-Aldrich. Sodium chloride, ethylenediaminetetraacetic acid (EDTA), hydrogen peroxide (H_2_O_2_), potassium dihydrogen phosphate, and disodium hydrogen phosphate were purchased from Merk, India. Polyvinylpyrrolidone (PVP) and Rhodamine-B (Rh-B) were bought from Sisco Research Laboratory Pvt. Ltd. (SRL), India. The L929 and 3T3 cells were procured from NCCS, Pune, India. Deionized water was used for all synthesis reactions and analysis of CDs.

### 2.1 Synthesis of CDs

The clove buds were washed thrice with distilled water to eliminate dust particles, followed by drying overnight at 100 °C in a hot air oven. The raw material was then ground into fine powder utilizing the mortar and pestle. Then, 0.5 g of fine clove powder with and without PVP was dispersed in 30 mL of water with continuous stirring and heating at 100 °C for 30 min using a mechanical stirrer (REMI, India). The resultant homogenous reaction mixture was transferred to a 60 ml capacity autoclave chamber (TECHNISTRO, India) at 200 °C for 5 h in the hot air oven. The larger particles in the supernatant were separated through centrifugation for 15 min at 10,000 rpm, followed by filtering through a 0.22-micron syringe filter (Sartorius, India). The filtrate containing PPCCDs was dried and reconstituted to a 10 mg/mL solution and stored at 4 °C for further experiments.

### 2.2 Characterization of CDs

CDs were systematically characterized to study their physicochemical properties that will further define their biological and non-biological applications. To analyze the functional groups, present in CDs Fourier Transform Infrared spectroscopy (FTIR) was performed by drop-casting and drying samples on KBr pellets using Nicolet 6700 (Thermo Fischer Scientific Instruments, USA). UV-Visible spectrometer 201 (Thermo Scientific, USA) was used to analyze the absorption spectra of synthesized samples. The fluorescent property was analyzed using a spectrofluorometer RF-6000 series (Shimadzu, India). The nanostructure of CDs was analyzed employing high-resolution transmission electron microscopy (JEM-2100 HR-TEM, JEOL, Japan) with 200kV accelerating voltage. In brief, the sample was drop cast on a carbon-coated copper grid with overnight drying at 25 °C to get rid of remnant moisture. The histogram for size distribution calculation representing the average particle size was plotted over 100 particle counts using Image J software. For elemental composition profiling of CDs, X-ray photoelectron spectroscopy (XPS, PHI 5000Verse Probe II (ULVAC-PHI Inc., Japan)) was performed.

### 2.3 Stability study of PPCCDs

The fluorescence intensity dependency on pH and UV light exposure are two key factors in investigating the stability of fluorescent dyes and also ensuring their applicability in biological environments. In brief, the aqueous solution of PPCCDs (0.05 mg/mL) was used with HCl (2 M) or NaOH (2 M) to adjust their pH, either acidic or alkaline. The same quantity of PPCCDs was also studied for the different exposure times of UV light (254 nm). The effect of pH variation and UV light exposure on fluorescence and absorption intensity was investigated using fluorescence and UV-Visible spectroscopy.

### 2.4 Antioxidant activity assay

#### 2.4.1 DPPH free radical assay

The *ex vivo* antioxidant efficacy of PPCCDs was evaluated by a DPPH assay with minor modifications [21]. The presence of an antioxidant molecule scavenges DPPH free radicals, reducing the purple-colored stable radical cation to a yellow-colored solution with a consequent decrease or loss of absorbance, which can be measured via a spectrophotometer.

In brief, 0.3 ml of methanolic solution of DPPH (0.5mM) was added to different concentrations of PPCCDs and incubated in the dark for 100 min. The optical density post-reaction completion was measured using a UV-Vis spectrophotometer at 517 nm. The experiment was repeated thrice with five batches of PPCCDs to ensure reproducibility. The antioxidant efficacy against the DPPH free radical was calculated using the following equation:

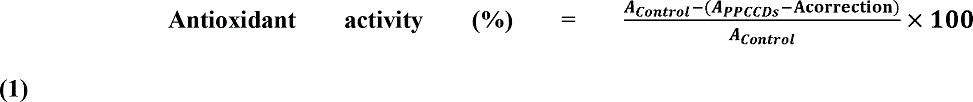

Here, A_correction_ is the absorbance of PPCCDs without DPPH and A_control_ is the absorbance without PPCCDs addition.

#### 2.4.2 Superoxide anion radical assay

The superoxide anion radical scavenging capability was investigated using the NBT assay with minor modification [22]. The reaction mixture of sodium carbonate (125mM, pH 10.2), NBT (24 µM), EDTA (0.1 mM), and hydroxylammonium chloride (1mM) without PPCCDs was taken as control. The reaction was initiated by adding different concentrations of PPCCDs to the above reaction mixture and recording the optical density at 560 nm after 30 minutes of incubation using a spectrophotometer. The superoxide inhibition percentage was calculated using the following equation.

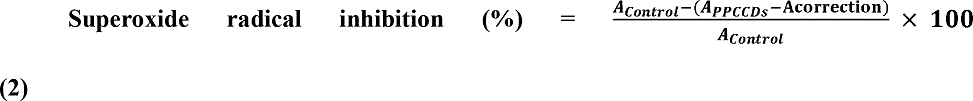

Here, A_correction_ is the absorbance of PPCCDs without NBT and A_control_ is the absorbance without PPCCDs addition.

### 2.5 Investigation of catalytic activity

The catalytic activity of PPCCDs was evaluated for the degradation of Rhodamine-B (Rh-B) in the presence of sodium borohydride (NaBH_4_) using the UV-Vis spectrophotometer. The reactants and catalyst concentration were established by following the modified protocol [6]. Briefly, two cuvettes each of 3.5 mL capacity were occupied with 0.20 mL of newly prepared NaBH_4_ (0.50 M), 1 mL of Rh-B (0.01mM), and 1.8 mL of water to make up a 3 mL complete solution. One of the above cuvettes was added with 20 μL (0.2 mg/mL) of PPCCDs, while the other served as a control for the study. The degradation of Rh-B was examined by capturing the time-dependent absorbance of the reaction mixture at 554 nm wavelength using UV-Visible spectroscopy after every 2 minute time interval. The degradation percentage (%) of Rh-B dye was calculated as per Eq. 3.

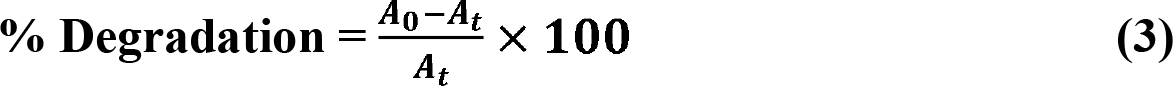

Where A_0_ is the initial absorbance at time t = 0, and A_t_ stands for the absorbance after reaction time t.

### 2.6 Evaluation of cytotoxicity

The toxicity of PPCCDs was evaluated using the MTT assay, wherein mitochondrial succinic dehydrogenase enzymes present in actively proliferating cells will reduce tetrazolium compound (MTT dye) into its insoluble purple color formazan crystal. In brief, 3T3 and L929 cells were seeded in 96 well plates at a cell density of 5000 cells/well in complete DMEM, high glucose media (Thermo Fischer Scientific, USA) containing Fetal bovine serum (10% FBS, Thermo Fischer Scientific, USA), and antimycotic-antibacterial (1%, Thermo Fischer Scientific, USA). A series of diluted concentrations of PPCCDs, i.e., 0.1, 0.3, 0.5, 0.7, 0.9, and 1.1 mg/mL were added to each well in triplicate post-48 h incubation in an incubator maintained at 37 °C, 5% CO_2_, and 95% humidity. After 48 h of treatment, the MTT reagent (5 μg/mL) was added in the dark environment and incubated for 4 h more. After incubation, an equivalent amount of DMSO was added to solubilize the purple formazan crystals, and the optical density was evaluated at 570 nm using the BIO-RAD microplate reader-550. The untreated cells were taken as a control for the study and processed similarly. The cell viability was evaluated using the following equation:

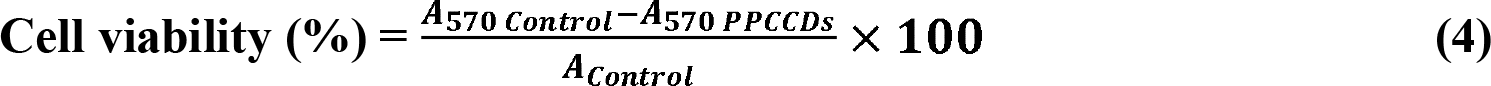

### 2.7 Live cell imaging

For assessing the bioimaging potential of PPCCDs, 3T3 and L929 cells were cultured on a poly-L-lysine-coated coverslip and incubated in a culture medium containing PPCCDs for 9 h. Subsequently, the cultured cells were repeatedly washed with PBS to avoid any overstaining before being captured under the inverted fluorescence microscope (eclipse Ts2R-FL, Nikon, Japan). The coverslip was imaged with the help of fluorescence microscopy in bright field and fluorescence mode.

## 3. RESULTS AND DISCUSSION

PPCCDs was successfully synthesized from clove buds using hydrothermal carbonization at 200°C for 5 h in presence of PVP as a passivating agent, as depicted in **Fig. S1**. The colors of non-passivated CDs (CCDs) and passivated PPCCDs were observed to be light yellow or dark greyish-black under white light respectively, while under UV light illumination, both CDs showed green fluorescence as shown in **Fig. S2**. The quantum yield of CCDs and PPCCDs were performed and calculated to be 7.6% and 10.9%, respectively, as shown in **Fig. S4**. This dictates the surface passivation of PVP results in improved quantum yield in comparison to non-passivated CDs (**Table S1**). The quantum yield and multifunctional aspects of various surface passivated CDs are compared in **Table S4**. The PPCCDs solution was darker when compared with the CCDs, which may be due to the passivation of CCDs owing to the addition of PVP.

### 3.1 Physiochemical characterization of CDs

#### 3.1.1 FTIR analysis

FTIR analysis of CCDs, PPCCDs, and pristine PVP (**Fig. 1a**) were studied for analysis of the functional groups and bonds associated with CDs. The stretching vibration of the O-H group and N-H group exhibit broadband in the range 3600-3200 cm^-1^ [23]. Two smaller peaks at 2922 cm^-1^ and 2862 cm^-1^ represent symmetric and asymmetric vibrations of the -CH_3_ and -CH_2_ groups, respectively. The presence of sp^2^ hybridized honeycomb lattice (aromatic C = C or C = O stretching) is indicated by a sharp peak at 1645 cm^-1^ in the case of pure PVP and PPCCDs, which have redshift for CCDs. The absorption peak at wavenumber 1287.6 cm^-1^ belongs to the C-N bond stretching in PVP [24]. The peak at 1050 cm^-1^ is ascribed to the stretching vibration bands of C-O on the PPCCDs surface, while the peak at 1027 cm^-1^ resembles the aromatic ether group present due to PVP addition [25]. The FTIR spectrum of PPCCDs represents the combination of various bond and functional groups present in the pristine PVP and CCDs spectrum collectively. The presence of functional groups at the surface of CDs makes them well dispersed in the aqueous medium.

**Fig. 1.**
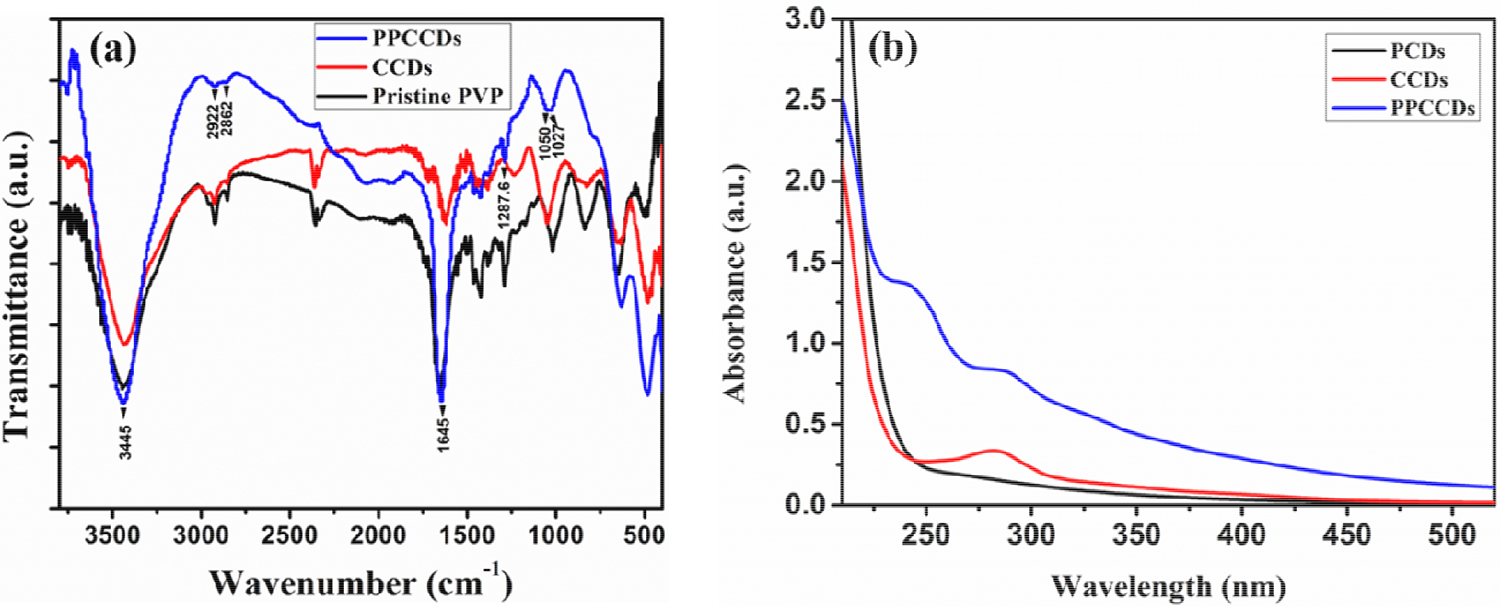
(a) FTIR spectra and (b) UV-Vis absorption spectra of passivated (PPCCDs) and non-passivated (CCDs) carbon dots along with FTIR spectrum of pristine PVP and UV-Vis spectrum of PVP-derived carbon dot (PCDs).

#### 3.1.2 Optical behavior of CDs

The photophysical analysis of non-passivated (CCDs), PVP-derived CDs (PCDs), and passivated CDs was performed to understand significant modulation in their optical properties on passivation, collectively. **Fig. 1b** indicates UV-Vis analysis of PPCCDs and CCDs with two broad absorption peaks at 288 nm and 282 nm, respectively, belonging to π–π* electron transition of C=C bonds [6]. While one more small shoulder peak at 324 nm in the PPCCDs sample resembles n–π* electron transition of C=O bonds [3,6]. For comparison, the UV-Vis spectrum of PCDs as shown in **Fig. 1b** indicates absorption in the deep ultraviolet region. The surface passivation is responsible for an increase in absorption intensity (in the wavelength range of 240-600 nm) of passivated CDs in contrast to the non-passivated CDs [26]. The comparative study of the fluorescent behavior of CCDs, PCDs, and PPCCDs as shown in **Fig. 2ace** revealed a significant difference in their fluorescent emission by varying the excitation wavelength in the range of 340-500 nm with a 20 nm increment. The passivated PPCCDs have three coexisting emission peaks centered at 415 nm, 439 nm, and 462 nm in comparison to non-passivated CCDs at 432 nm, which attained their maxima when excited at 380 nm and 360 nm excitation wavelengths, respectively. While PCDs is showing maximum emission at 417 nm when excited by 340 nm wavelength. Specifically, the fluorescence peak at 432 nm for CCDs and 417 nm for PCDs was shifted at 439 nm and 415 nm, respectively, in the case of PPCCDs. Notably, an additional peak at 462 nm has appeared in the case of PPCCDs as a result of passivation that might be due to the synergistic effect. Pal et al. also reported a similar result for CDs derived from curcumin while passivated by polyethyleneimine [14]. They have shown the synergistic effect on the fluorescence behavior of passivated CDs as compared to the CDs derived from their native ones. As shown in **Fig 2bdf**, samples were excited within the wavelength range of 340 nm to 500 nm, PCDs had emission at a regular interval, while CCDs had relatively different phenomena. Initially, peaks of emission spectra at the excitation wavelength range of 340 to 380 nm had marginal differences while spectra at 380 nm and above a relatively higher magnitude of peak shift was observed. Interestingly, this clustering effect of emission spectra for CCDs was further evident in the case of PPCCDs as a result of passivation with PVP. The multiple emission peaks of PPCCDs show both types of fluorescent characteristics, such as excitation independent emission at and below 400 nm wavelength and excitation dependent emission above 400 nm excitation wavelength, similar to reported literature [27,28]. The unique uniform surface emissive trap of passivated CDs at lower wavelengths is valuable for avoiding autofluorescence issues and is also responsible for excitation independent emission [27]. While the excitation wavelength-dependent emission behavior of CDs is an outcome of the various surface emissive traps with different energy levels at higher wavelengths [28]. One possible explanation for coexisting peaks is the occurrence of different electron transitions inside the graphitic core as a result of different size effects and surface defects in passivated CDs [29,30]. The photophysical properties of PPCCDs are more efficient and improved than CCDs and were further studied and explored for multiple applications.

**Fig.2.**
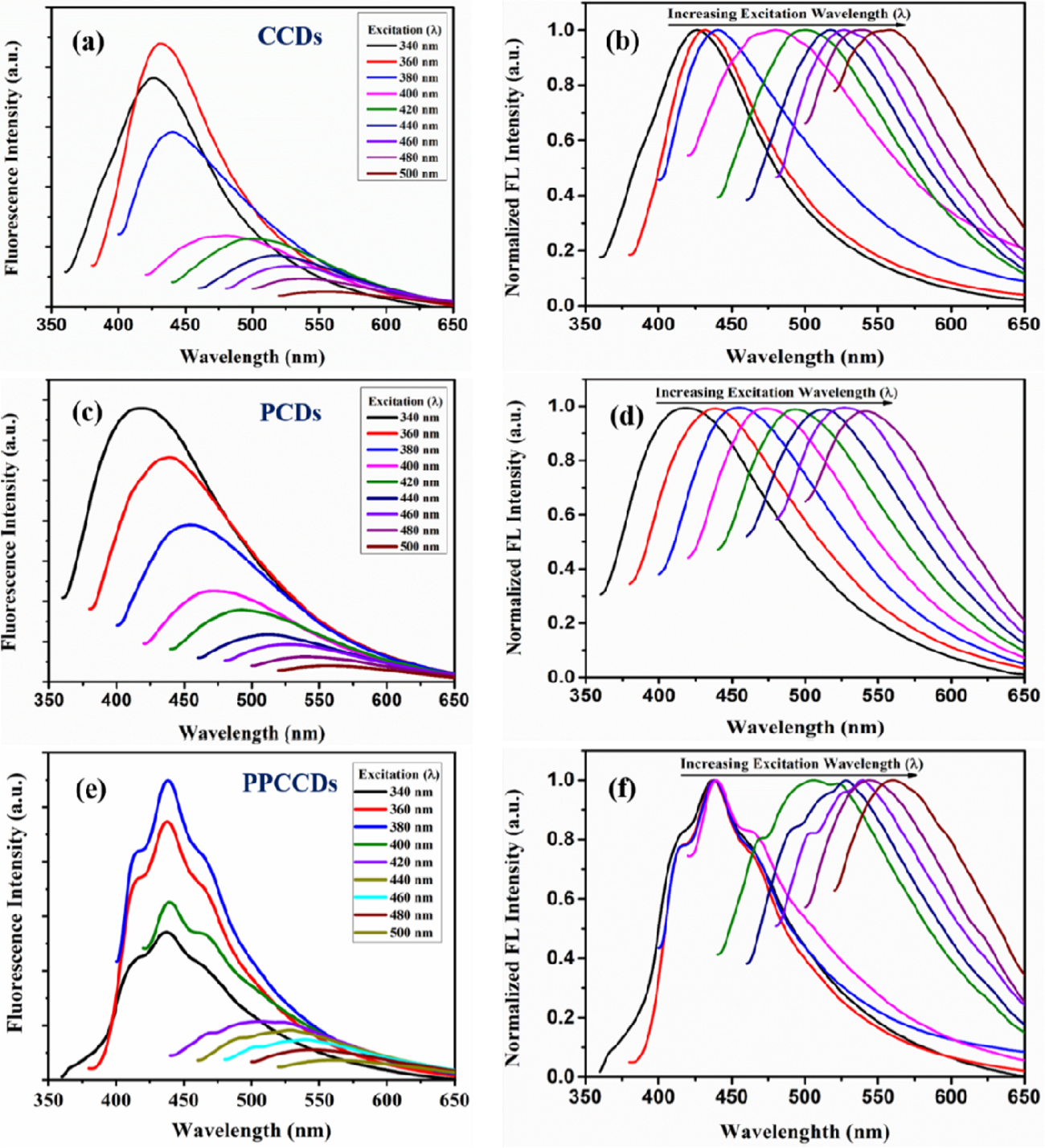
(a, c, and e) representing fluorescent emission spectra, and (b, d, and f) showing normalized fluorescent spectra of CCDs, PCDs, and PPCCDs by varying the excitation wavelength in the range of 340-500 nm.

#### 3.1.3 Morphological, Structural, and Chemical characterization

The structural and morphological analysis shown in **Fig. 3** represents the XRD data, TEM image, size distribution histogram, and HRTEM image of PPCCDs. The XRD analysis reveals one broad peak at 2θ = 23.54°, which corresponds to the (002) plane of graphitic carbon of crystalline nature [31]. In the case of CCDs, the XRD peak is a little shifted to 24.4° as shown in **Fig S5a**. While TEM images demonstrate the spherical shape of PPCCDs with an average diameter of 2 nm. The histogram plot was obtained by using image-J software, which resembled different-sized particles with size distribution window in the range of 1-5 nm. The HRTEM image showed lattice fringes with a d spacing value of around 0.33 nm, indicating the presence of a (002) plane of the graphitic carbon validated with XRD data. The TEM analysis of CCDs showed the presence of agglomerated CDs, as shown in **Fig S5b,** that was mostly stabilized after passivation.

**Fig. 3.**
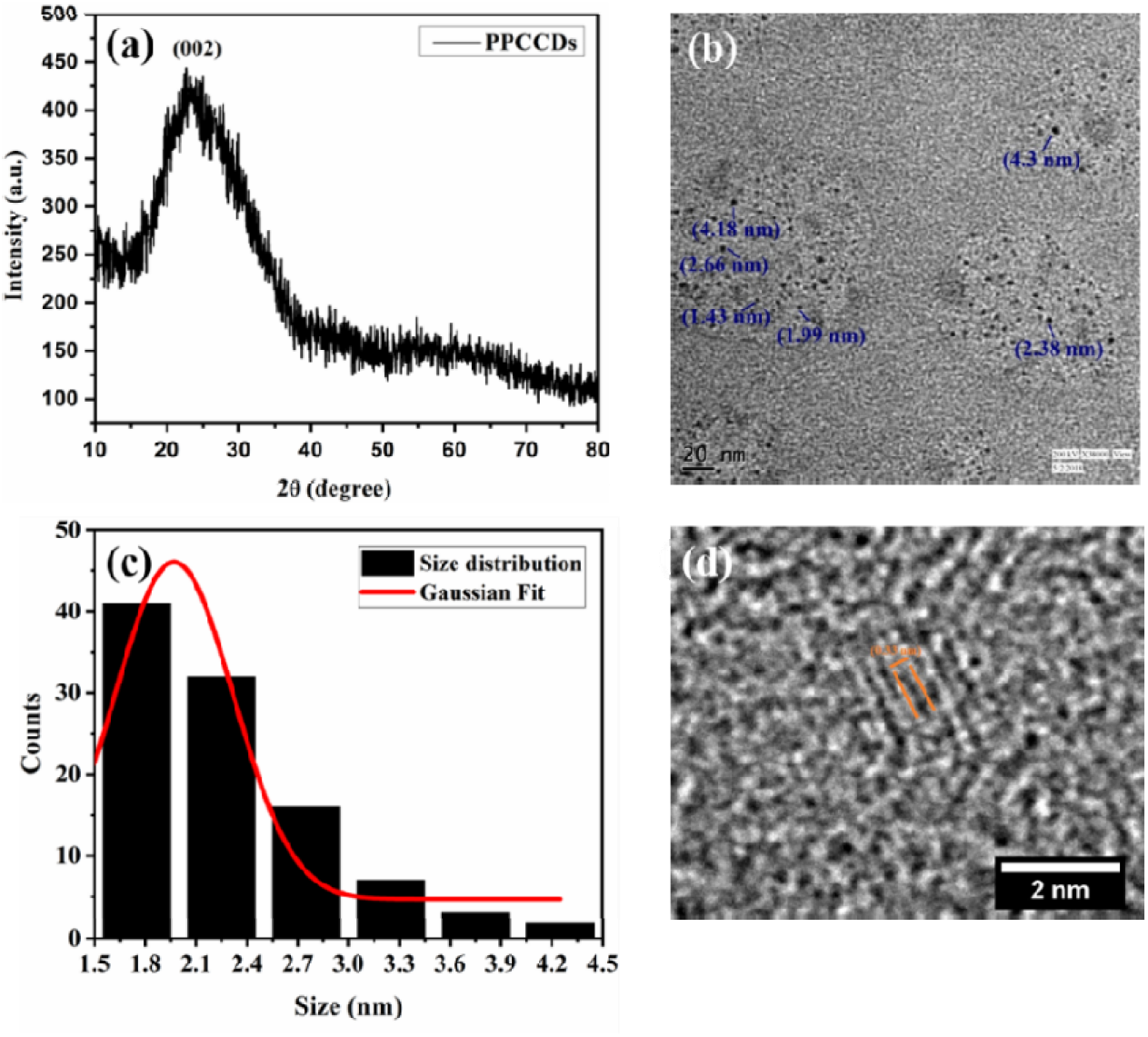
(a) XRD spectrum, (b) TEM image, (c) respective size distribution histogram, and (d) HRTEM fringe pattern image of PPCCDs.

X-ray photoelectron spectroscopy (XPS) was performed to further examine the surface chemical state and composition of PPCCDs. The clove buds mostly contain polyphenolic components dictating the presence of hydrogen, carbon, and oxygen elements in their chemical structure [32,33]. The full scan spectrum as shown in **Fig. 4a** revealed the existence of carbon, oxygen, and nitrogen, atoms with an atomic ratio of 85.30%, 10.50%, and 4.20%, respectively.

**Fig. 4.**
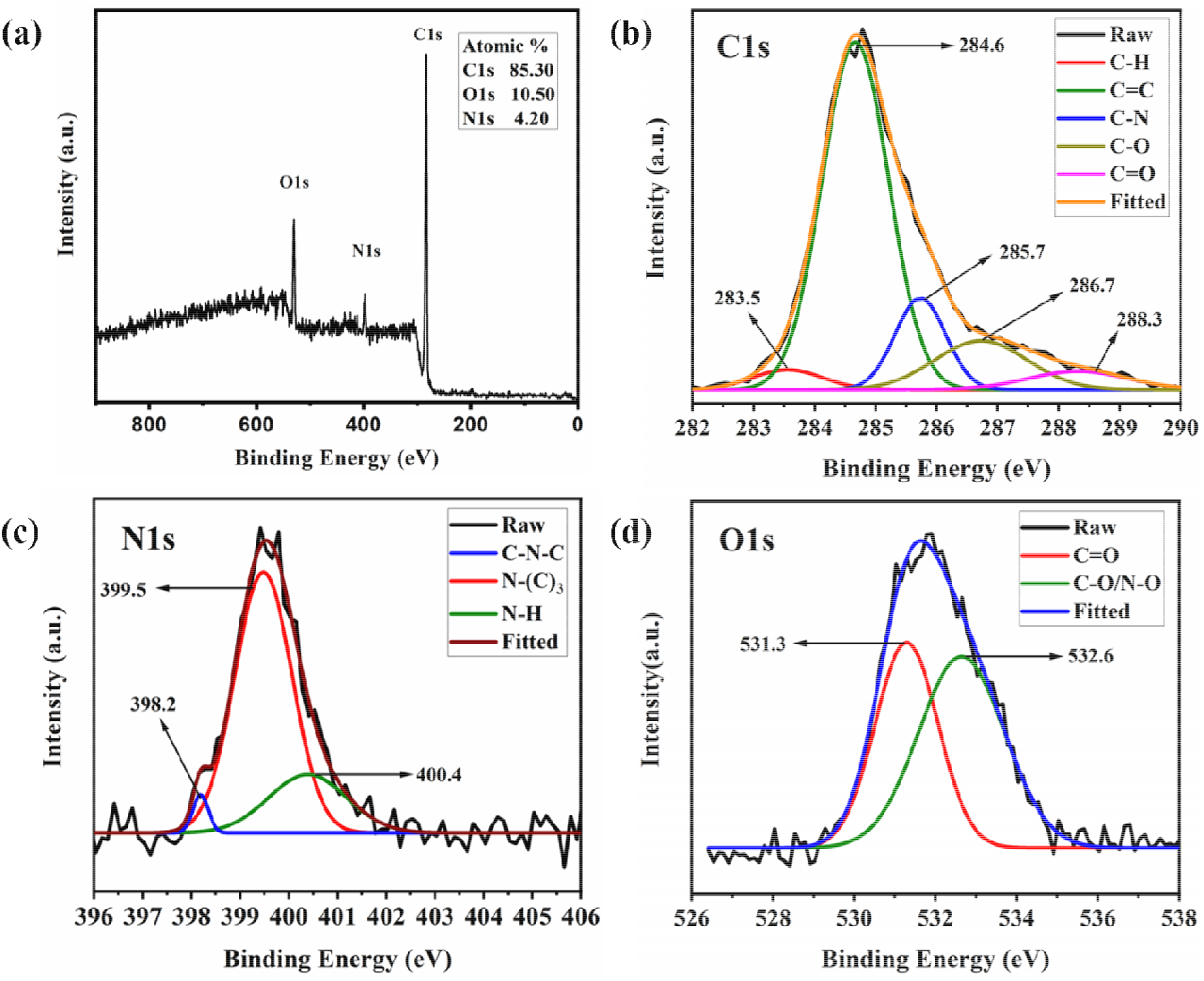
(a) The full scan XPS spectrum and HR-XPS spectrum of (b) C1_s_, (c) N1_s_, and (d) O1_s_ of synthesized carbon dots (PPCCDs).

High-resolution spectra of C1_s_, N1_s_, and O1_s_ as shown in **Fig. 4b-d** dictated the presence of various atomic bonds validation with FTIR data. The deconvoluted C1_s_ spectrum revealed the presence of C-H, C=C, C-N, C-O, and C=O bonds with their respective binding energies of 283.5, 284.6 eV, 285.7 eV, 286.7 eV, and 288.3 eV [14,23]. Furthermore, the deconvoluted spectrum of the N1_s_ peak shown in **Fig. 4c** revealed three peaks at 398.2 eV, 399.5 eV, and 400.4 eV that are associated with the functionality of the C-N-C, pyrrolic N, and N-H bonds, correspondingly, confirm the presence of nitrogen doping in CDs [14,31]. **Fig. 4d** revealed deconvoluted O1_s_ spectrum with two peaks at 531.3 eV and 532.6 eV belonging to C=O, C-O/N-O bonds, respectively [34].

### 3.2 Stability of PPCCDs

The effect of pH and ultraviolet light irradiation on PPCCDs was performed to establish their stability under the studied conditions. As shown in **Fig. 5a**, pH variation does not have any significant change in the absorption intensity and spectra of PPCCDs, revealing the absence of aggregation in the studied conditions. The fluorescence spectrum was also observed with a corresponding change in pH values. Interestingly, at pH 7 the fluorescence spectra showed a similar intense band equivalent to the intensity at pH 11 supporting its suitable applicability for live-cell imaging as shown in **Fig. 5b**. Further, the impact of bleaching on PPCCDs has also been assessed post UV exposer over 3 h. As shown in **Fig. 5c-d**, there was no significant change in peaks intensity or position for both absorption and fluorescence spectra, revealing post UV exposure PPCCDs neither aggregated nor lost their intensity significantly beneath understudied conditions.

**Fig. 5.**
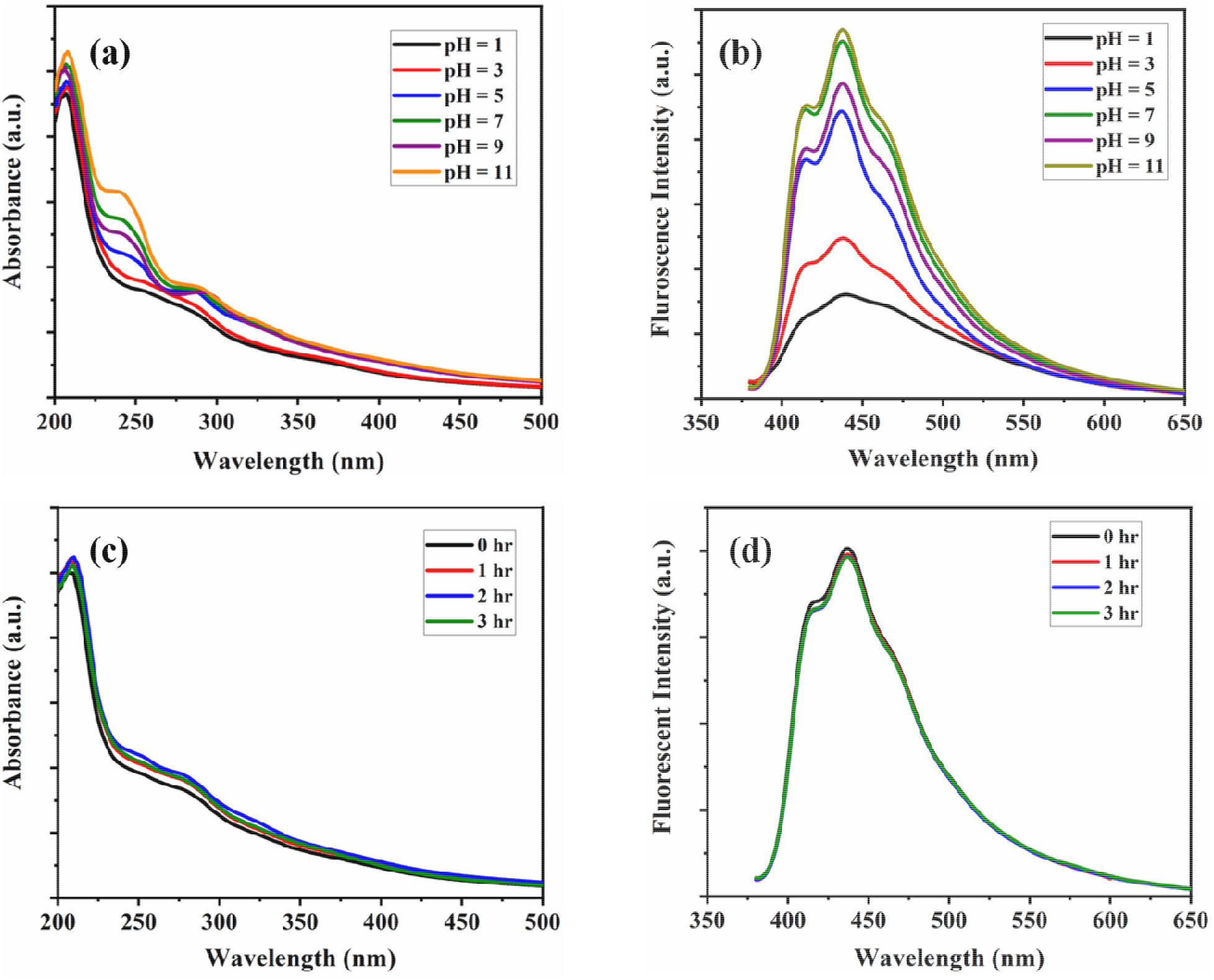
(a) and (c) represent UV-Visible spectra, (b) and (d) showing the fluorescent spectra of variant pH (in range 1-11) and ultraviolet exposer (λ = 254 nm) of the PPCCDs sample: Excitation wavelength for fluorescence emission was 360 nm.

### 3.3 DPPH free radical scavenging activity

The major problem regarding the generation of reactive oxygen species or free radicals in biological systems could be scavenged or neutralized by nanomaterials having good antioxidant activity [21,35–38]. The CDs have both electron donors and acceptors capability that makes them act as good antioxidant and prooxidant agents [39–41] The CDs are a less explored and more suitable candidate for antioxidant study due to their minimal toxicity [3,4,31].

The antioxidant efficacy of PPCCDs was measured by using a standard DPPH free radical assay. It was noticeably detected from **Fig. 6a** that the absorption peak of DPPH radicals at 517 nm is decreased when the concentration of PPCCDs increased from 15 to 500 µg/mL in the reaction mixture. **Fig. 6b** represents the dose-dependent scavenging ability of PPCCDs which indicated an increase in antioxidant activity percentage attained their maximum of 94% at the highest concentration of 500 µg/mL. And the EC_50_ (concentration required to reduce 50% of the original DPPH free radical) value of PPCCDs was calculated to be 57 µg/mL. The lower EC_50_ value of PPCCDs signifies that it is more potent at scavenging DPPH free radicals than previously reported CDs [4,31,42]. The antioxidant efficacy of PPCCDs toward DPPH radical degradation was found to be superior in comparison to other reported CDs at their highest reported concentrations as shown in **Table S2**.

**Fig. 6.**
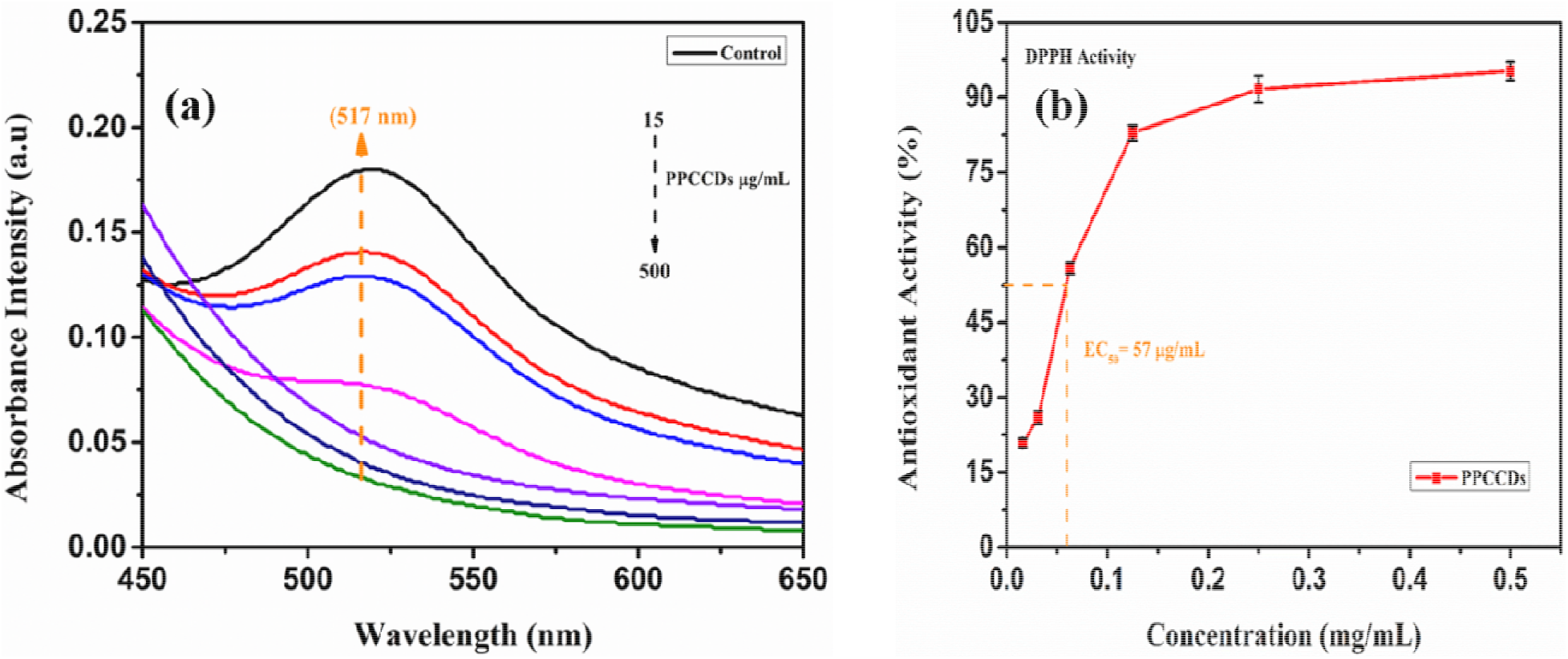
(a) DPPH reduction absorption spectra and (b DPPH radical scavenging activity plotted against increasing PPCCD concentrations with their EC_50_ value.

#### 2.4.2 Superoxide anion radical scavenging activity

The superoxide radical scavenging activity of PPCCDs was performed using an NBT assay. The mechanism involves the generation of superoxide anion by auto-oxidation of hydroxylammonium chloride while maintaining sodium carbonate at basic pH of 10.2. The generated superoxide anion in the presence of a catalyst inhibits the reduction of NBT. The UV-Vis spectrum as shown in **Fig. 7a** indicated an increase in superoxide anion radical scavenging potential of PPCCDs on varying concentrations from 31 to 500 µg/mL. It could also be observed that at a higher concentration of PPCCDs more absorption intensity is observed toward the ultraviolet region. The inset of **Fig 7a** showed a decrease in absorption intensity at 560 nm on increasing the concentrations of PPCCDs. **Fig. 7b** represents the percentage of superoxide radical inhibition with a change in concentrations, indicating the SOD-mimicking ability of PPCCDs. The superoxide radical inhibition of 89% acquired at a concentration of 500 µg/mL along with an EC_50_ value of 53 µg/mL calculated from **Fig 7b** is comparable or better than other previously reported literature shown in **Table S2** [4,21,43].

**Fig.7.**
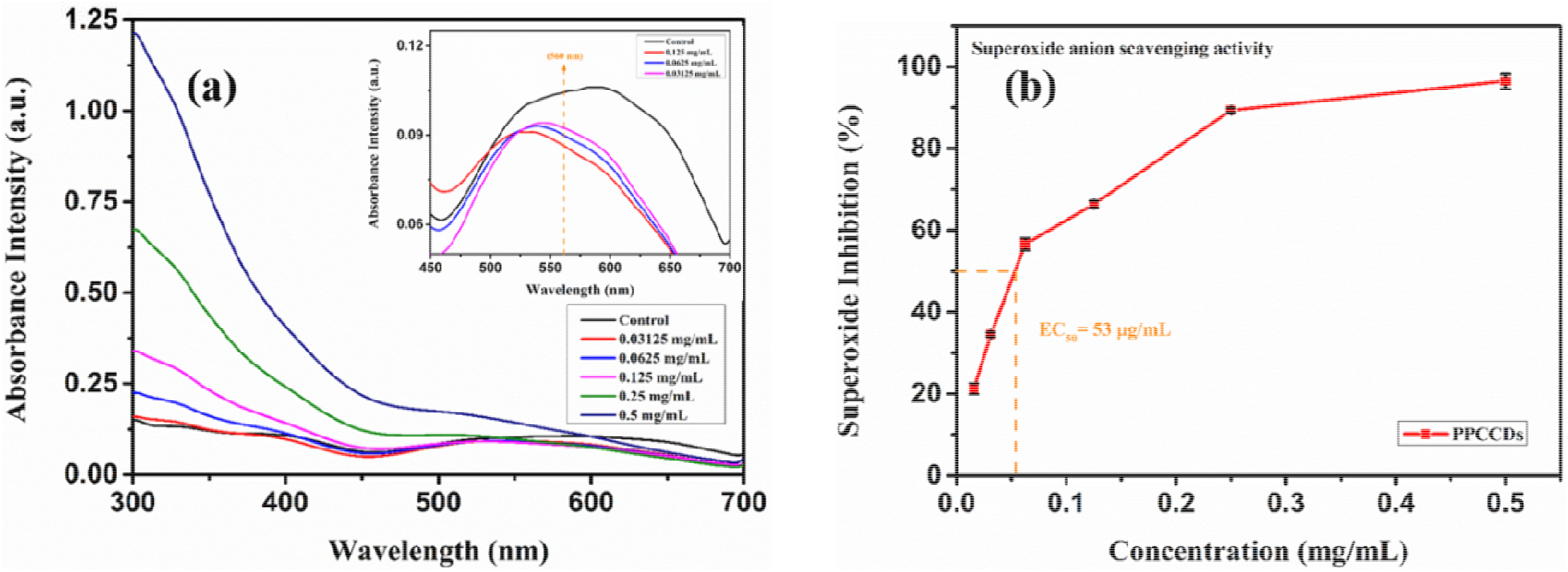
(a) NBT reduction inhibition absorption spectra (Inset shows magnified absorption spectrum at 560 nm.) and (b) superoxide radical inhibition activity plotted against increasing PPCCD concentrations.

### 3.4 Catalytic activity over Rh-B

In this study, Rh-B as a modal pollutant was designated to inspect the catalytic activity of PPCCDs using NaBH_4_ as a reducing agent. Rh-B is a commercialized organic dye most efficiently used in the textile, leather, and biological industry. It harms the environment and human health when disposed of in river water as waste without pre-treatment. Its accumulation in water results in various neurotoxic, mutagenic, and reproductive health issues in both animals and humans [44]. So, the demand for simple and cost-effective techniques along with efficient and non-toxic catalyst material is desirable for the removal of organic dyes from wastewater.

The aqueous solution of Rh-B shows a pink-red color with a characterized maximum absorption peak at a wavelength of 554 nm. **Fig. 8** represents the reduction of Rh-B using NaBH_4_ with or without the addition of PPCCDs as a catalyst for 20 minutes along with their degradation and rate kinetics. It can be demonstrated from **Fig. 8a-b** that the peak intensity of Rh-B dye decreased more gradually with time after adding NaBH_4_ alone in comparison to adding NaBH_4_ in the presence of PPCCDs. Hence, PPCCDs addition to the reaction mixture stimulated the degradation of dye as observed by a significant reduction in absorbance intensity. The plot of reaction kinetics of dye degradation with or without catalysts was evaluated by monitoring the change in absorbance concerning reaction time. The dye degradation kinetics were studied and found to have a first-order linear relationship, as described by the equation:

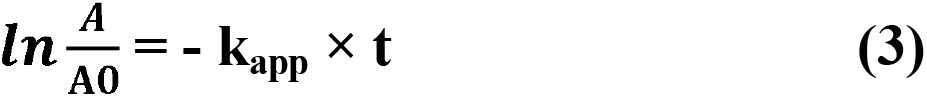

**Fig. 8.**
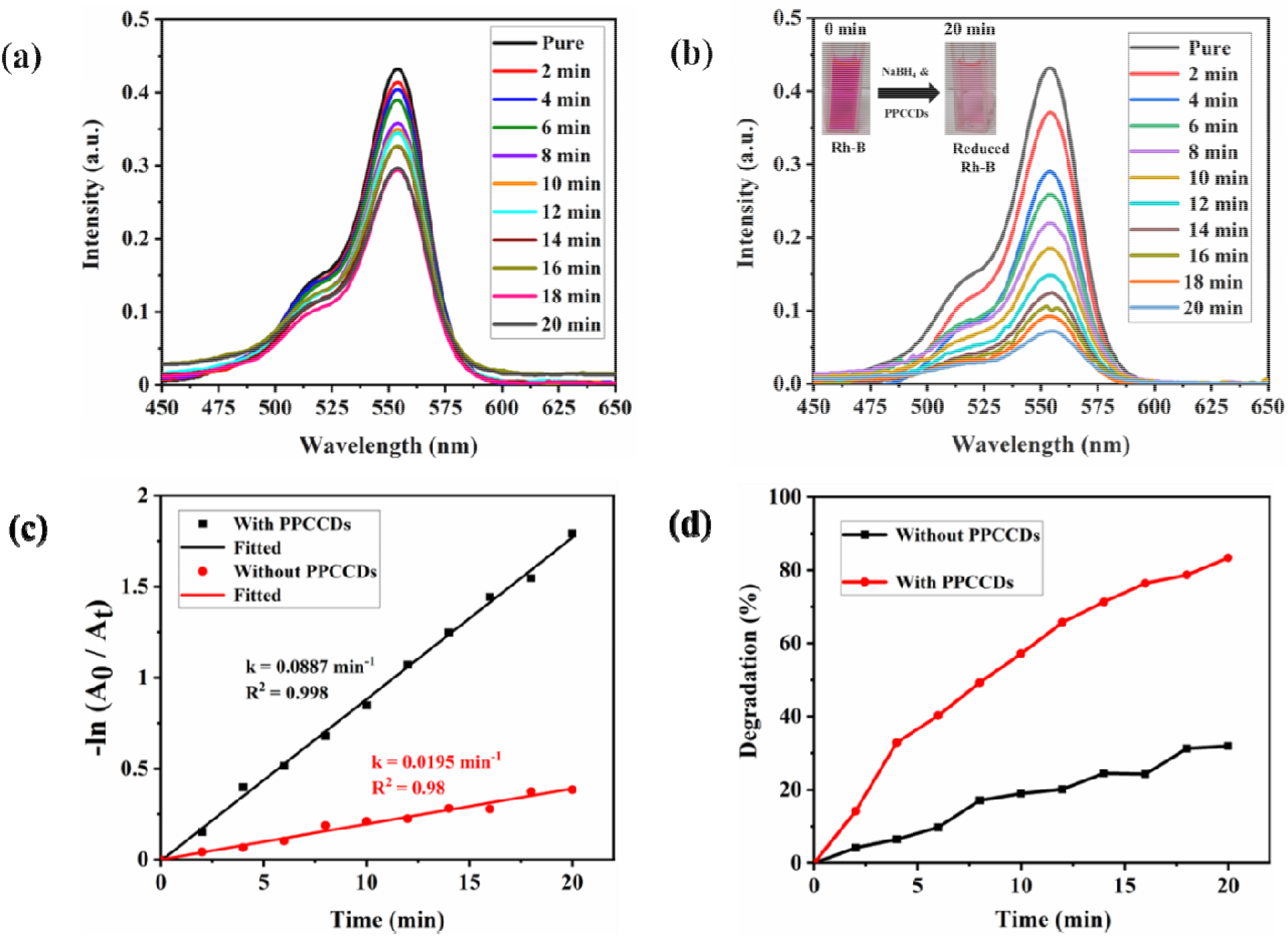
(a) and (b) showing reduction of Rh-B captured by UV-Visible spectroscopy, (c) reaction kinetics plot, and (d) representing degradation plot of Rh-B dye using NaBH_4_ with/without adding PPCCDs. Inset in Fig. 9b shows decolorization of Rh-B in the presence of PPCCDs as a catalyst.

Here, A_0_ and A are the absorbance at times t = 0 and t, k_app_ = apparent rate constant, and t = reaction time.

The linear dependency of ln (A/A_0_) concerning time as shown in **Fig. 8c** confirms first-order rate kinetics of the reaction with an apparent rate constant of 0.0887 and 0.0195 min^-1^ with R^2^ values of 0.998 and 0.98 in the presence and absence of catalyst, respectively. The specific rate constant was calculated by dividing the apparent rate constant k_app_ with the mass of catalyst (m) used, k^’^= k_app_/m in mg. The specific rate constant of PPCCDs was calculated to be 7.39 s^-1^ g^-1^. The specific rate constant k and apparent rate constant k_app_ of PPCCDs was found to be superior when compared with some other reported catalysts shown in **Table S3** [45–49]. The presence of PPCCDs results in faster degradation of Rh-B dye by nearly 86% (**Fig. 8d**) within 20 minutes, showing that the synthesized PPCCDs have good catalytic activity in the studied conditions.

The CDs due to their small size and high surface-to-volume ratio could be projected as an effective substrate for dye degradation. The process of degradation of Rh-B dye using PPCCDs as catalysts in the presence of NaBH_4_ can be better understood by the Langmuir-Hinshelwood mechanism represented in **Fig. 9**.

**Fig. 9.**
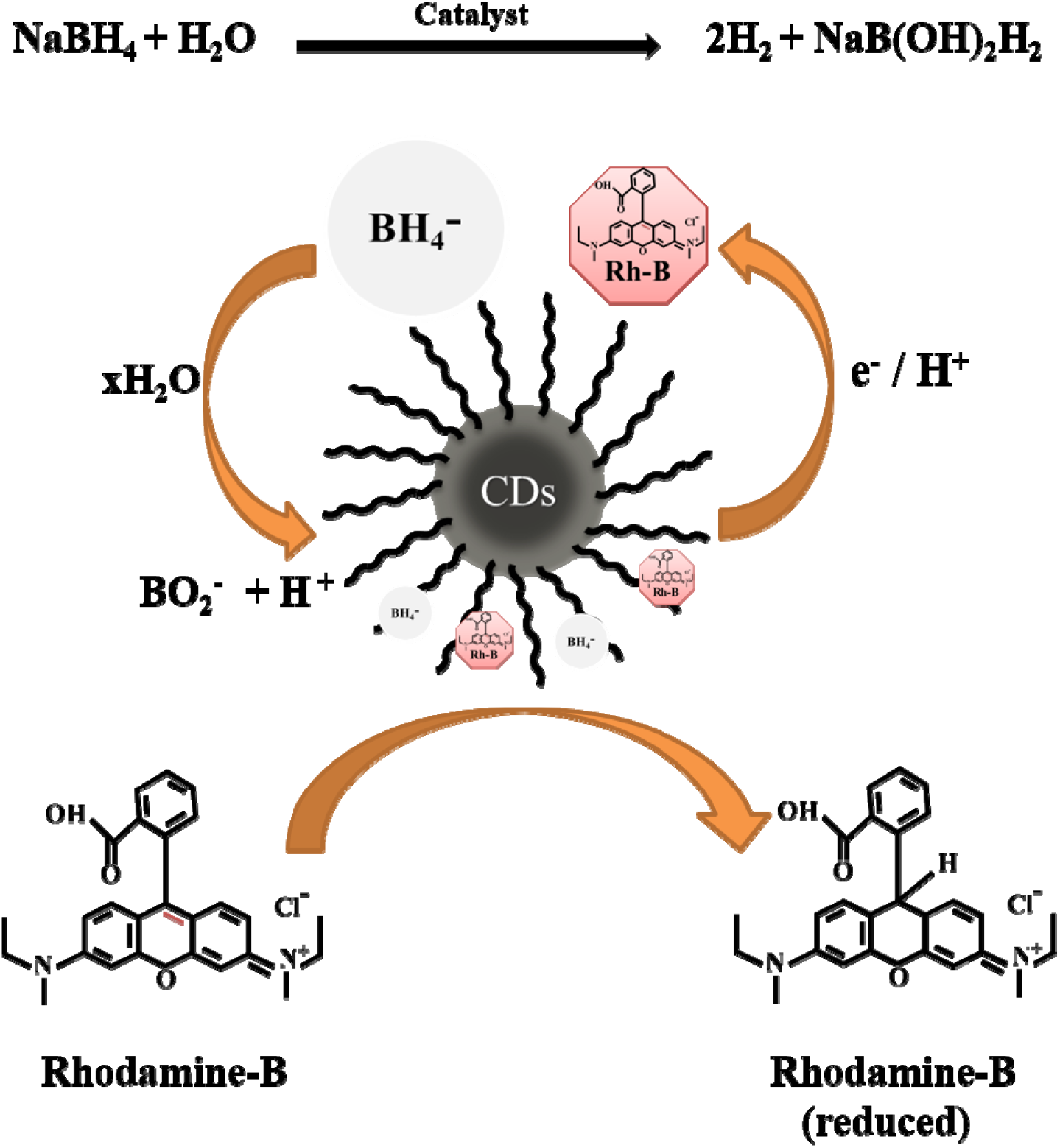
Schematic of the chemical reaction and process mechanism of Rh-B dye degradation using NaBH_4_ as a reducing agent and carbon dots as a catalyst.

The reaction mechanism involves the oxidation of an aqueous solution of sodium borohydride in the presence of CDs as a catalyst to liberate hydrogen bubbles [50]. The borate ions (BH_4_-) adsorb over the surface of catalysts oxidized with water molecules to generate BO_2_-ions and protonated hydrogen atoms. The role of the catalyst here is to decrease the reductive potential of NaBH_4_ and increase the reductive potential of Rh-B dye due to their nucleophilic and electrophilic nature. Further, the Rh-B dye and borate ions (BH_4_-) are adsorbed over the surface of CDs acting as a moderator (electron relay system) to receive an electron from borate ions and hand over the electron back to Rh-B through its surface [6]. Breaking of the conjugated system in the molecule and decrease of the potential barrier between reactant and product by evolving continuous H_2_ gas bubbles during this reduction reaction is explained in **Fig. 9**. The pink colors of Rh-B decolorized to colorless due to the destruction of the dye chromophore structure into less toxic species. Hence, the large surface-to-volume ratio due to smaller size and efficient electron-accepting potential due to nitrogen doping might be reasons which are credited with the good catalytic activity of PPCCDs.

### 3.5 In-vitro Cytotoxicity Test

Low cytotoxicity is one of the most important requirements for bio-imaging agents and a major drawback of semiconductor QDs employed in biomedical areas. Hence, to recognize the potential cytotoxicity of the PPCCDs, an MTT assay was performed using 3T3 and L929 cells. **Fig. 10** showed that the cell viability of PPCCDs for both cell lines was more than 80% after 48 h of treatment with a PPCCDs solution at the concentration of 0.5 mg/ml, and a remarkable decrease in cell viability was observed when higher doses of PPCCDs were employed. These observations signify that PPCCDs is a feasible candidate for cellular imaging in biomedical applications when used in appropriate concentrations.

**Fig. 10.**
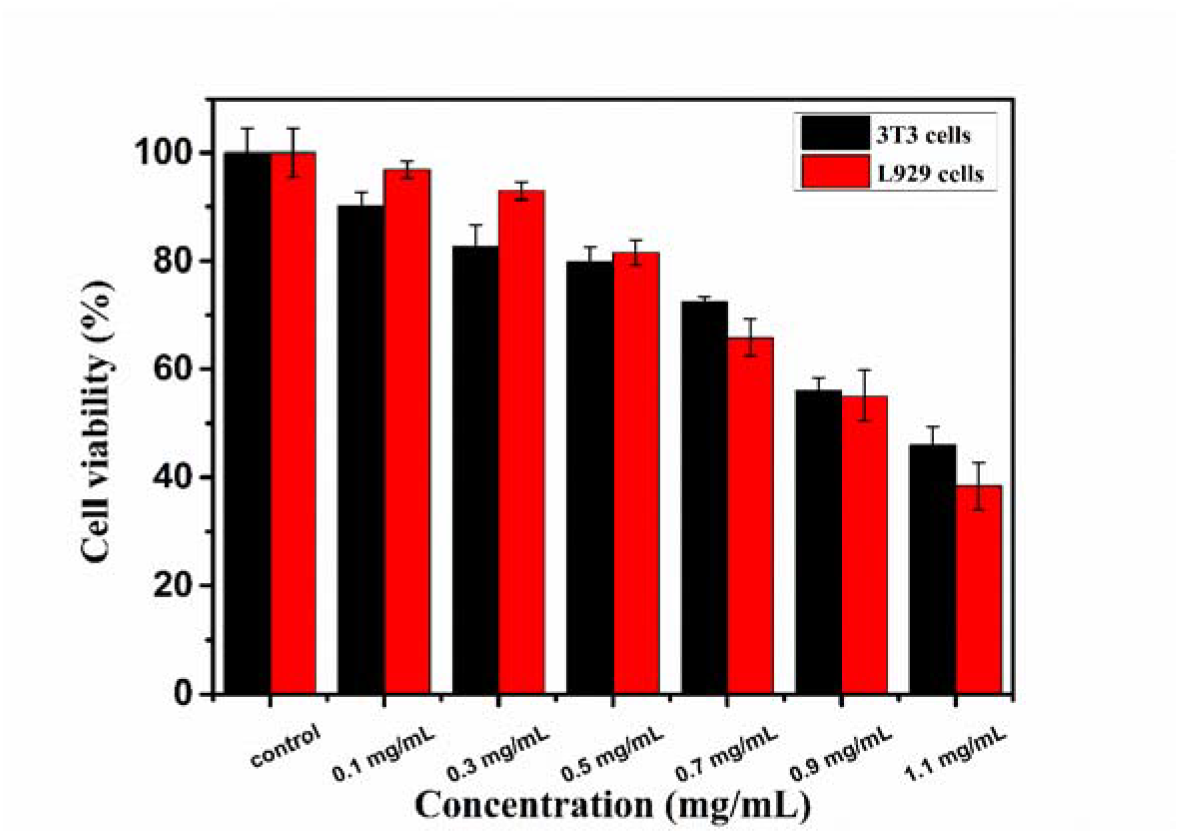
Cytotoxic evaluation of prepared concentrations of PPCCDs using MTT assay on 3T3 and L929 cells after 48 h incubation.

### 3.6 Live-cell imaging experiment

The bright-field and respective multicolor fluorescent images of 3T3 and L929 cells treated with PPCCDs at the different excitation wavelengths of 405 (blue), 488 (green), and 542 nm (red) were captured using a fluorescence microscope as shown in **Fig. 11**. The close investigation revealed that PPCCDs was more efficiently taken up by the cell membrane and cytoplasm region, as evident by the strong fluorescence. The possible mechanism for the entry of PPCCDs inside the cell could be due to endocytosis. The untreated L929 cells (without PPCCDs labeling) as shown in **Fig. S3** was found to be non-emissive when excited by lasers of wavelength 405 nm, 488 nm, and 542 nm, respectively. The findings are consistent with previous studies in which CDs were shown to stain cytoplasm and cell membrane [31,51,52]. The multicolor imaging capability of PPCCDs is better than various previously reported CDs [3,26,53,54]. This signifies that the synthesized PPCCDs could serve as an efficient bioimaging tool.

**Fig. 11.**
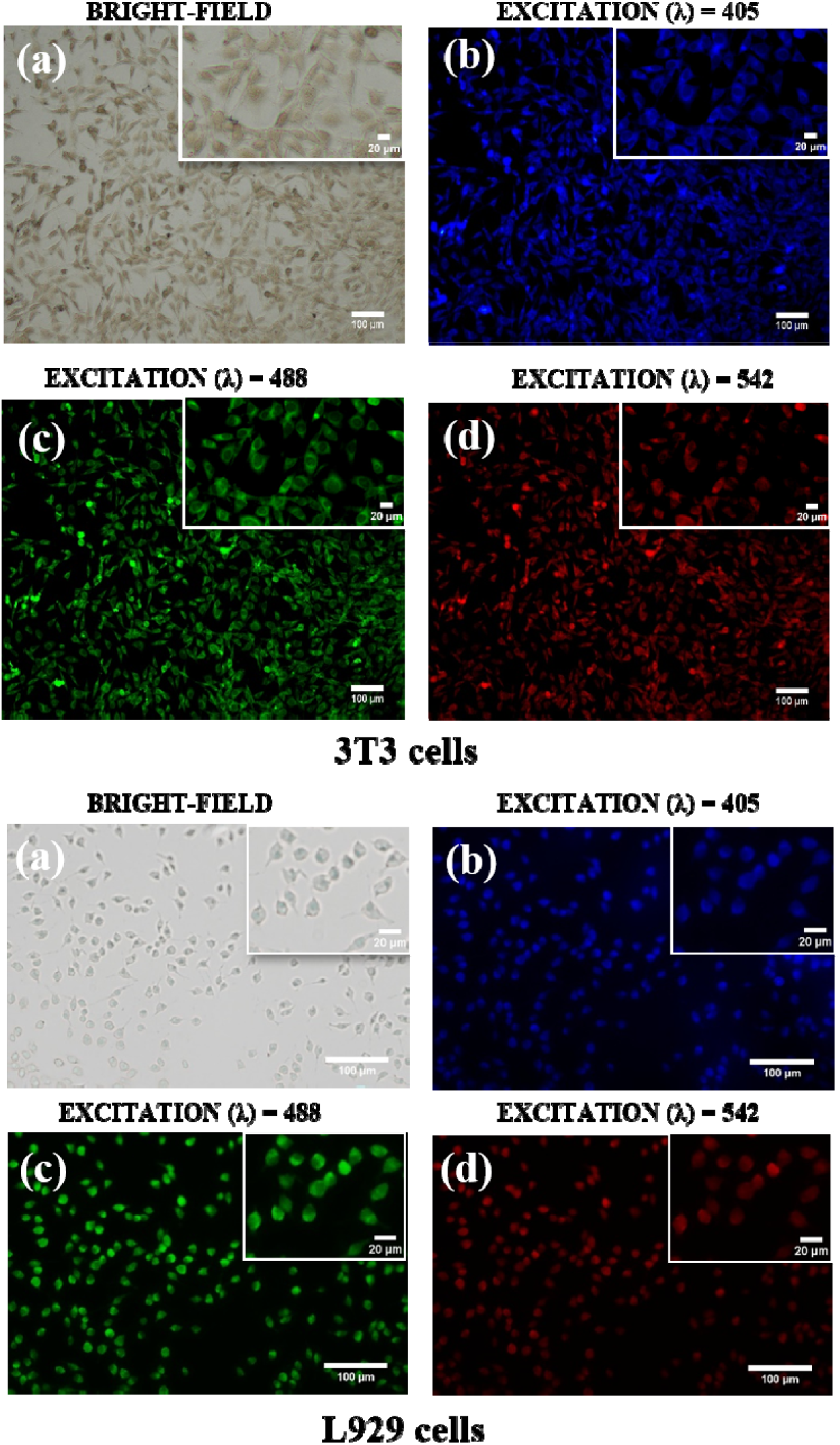
The fluorescent microscopy images representing in (a) bright-field and (b-d) fluorescent imaging of 3T and L929 cells incubated with synthesized PPCCDs for 9 h. The fluorescent images were captured at an excitation wavelength of (b) 405nm, (c) 488nm, and (d) 542nm. Scale bar: 100 µm. Inset shows magnified images at Scale bar: 20 µm.

## 4. CONCLUSION

The one-step synthesis of surface passivated clove buds-derived CDs via the hydrothermal method has been reported in this article. The optical studies were performed over CCDs, PCDs, and PPCCDs to understand the effects of passivation on their photophysical properties. The PPCCDs have excitation independent emission at a lower wavelength that is pivotal to avoid autofluorescence. Structural and morphological studies were performed to confirm the formation of spherical and graphitic CDs with a diameter of ∼ 2 nm. The photostability and pH stability tests were performed that demonstrated the insignificant effect of UV exposure on PPCCDs emission for biological applications. The cytocompatibility test of PPCCDs has shown more than 80% cell viability up to concentrations of 500 µg/mL using both cell lines making them a potential candidate for bioimaging. PPCCDs was effectively taken up by 3T3 and L929 cells and confined mainly to the cytoplasm and cell membrane regions, highlighting their potential as fluorescence imaging nanoprobes. Synthesized PPCCDs demonstrated excellent DPPH scavenging activity of 94% and superoxide inhibition activity of 89% at the highest concentration of 500 µg/mL along with a lower EC_50_ value of 57 µg/mL and 53 µg/mL. PPCCDs showed prominent catalytic activity in the degradation of the organic dye Rh-B using NaBH_4_. Overall, PPCCDs was synthesized by an environmentally friendly process to serve as an efficient bioimaging tool, an excellent antioxidant, and a good catalyst.

## Acknowledgment

The authors would like to thank MHRD (Ministry of human resource development), India for fellowship support and CRF (Central research facility), IIT Kharagpur for physiochemical characterizations. The authors are grateful to SNST, SMST, the Department of Physics, and the Department of Chemistry for their valuable contributions to the analysis.

