## Supplementary data for "Hydrothermal Synthesis of Clove Bud-Derived Multifunctional Carbon Dots Passivated with PVP- Antioxidant, Catalysis, and Cellular Imaging Applications"

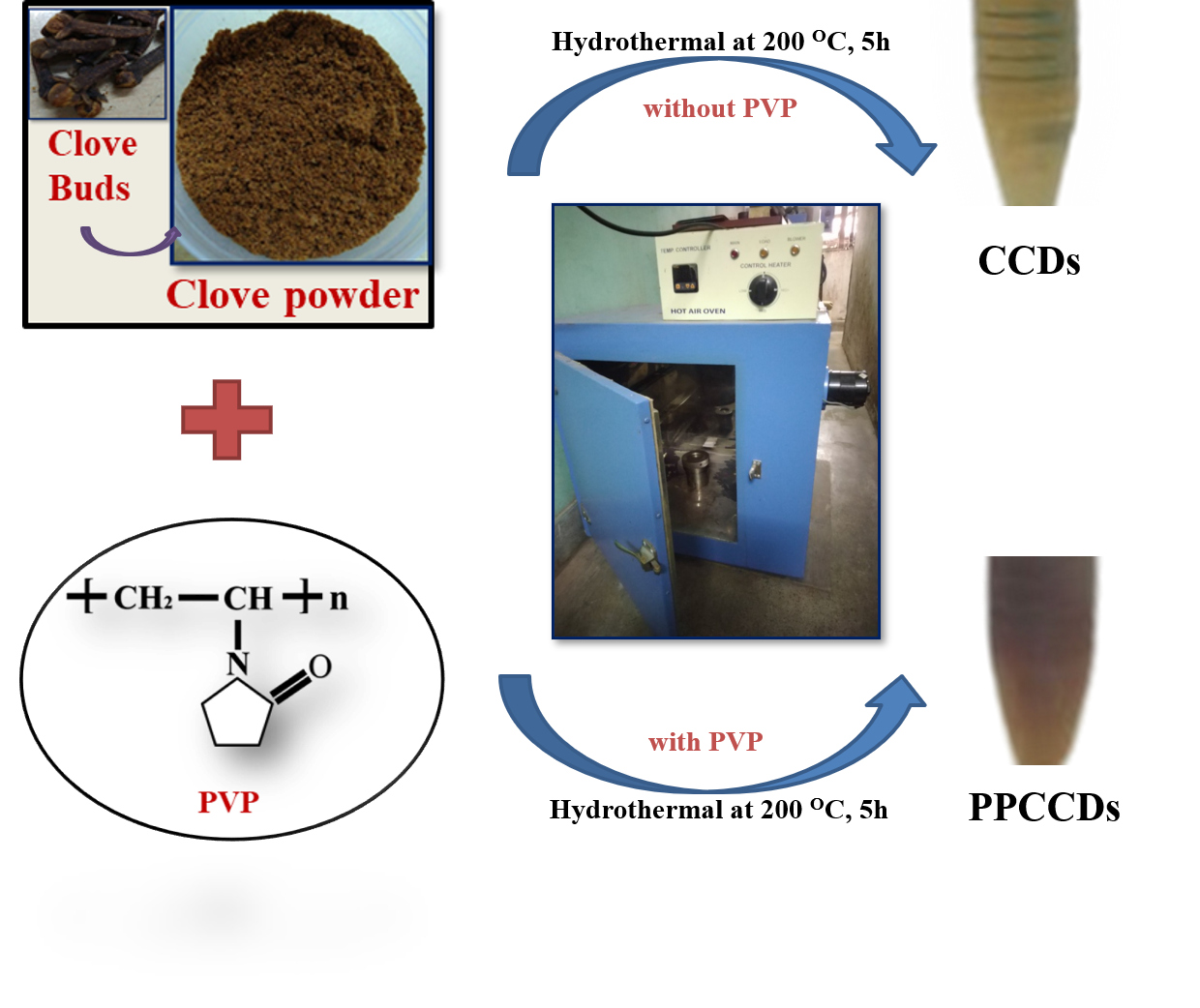

**Fig. S.1** Schematic of process involve in synthesis of PVP-passivated clove-derived carbon dots (PPCCDs) and non-passivated clove-derived carbon dots (CCDs)

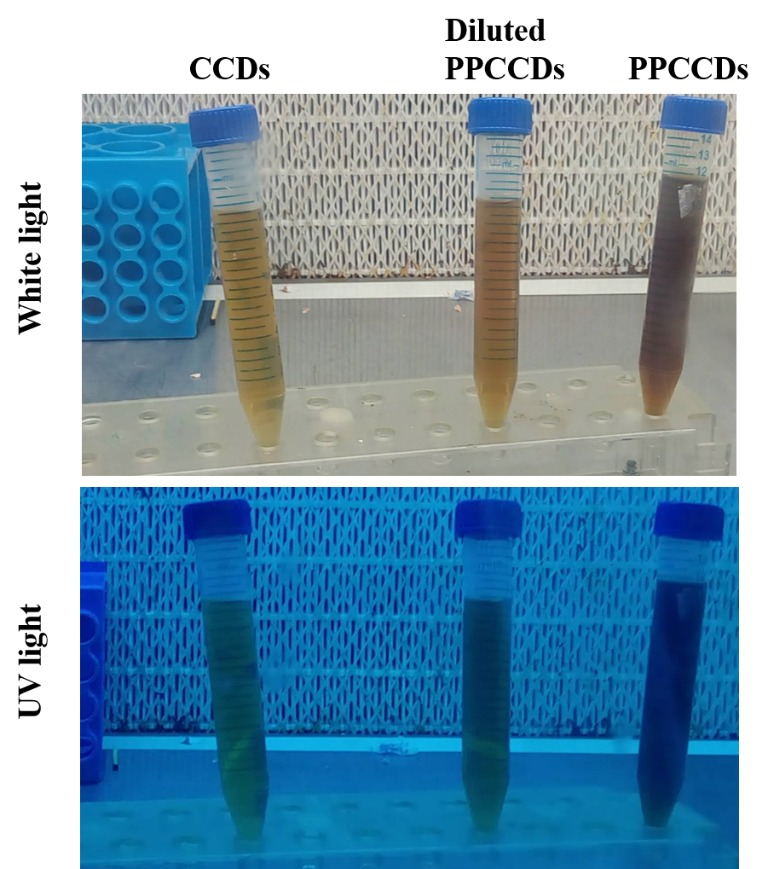

**Fig. S.2** White light and ultraviolet light exposer of as-synthesized PPCCDs and CCDs.

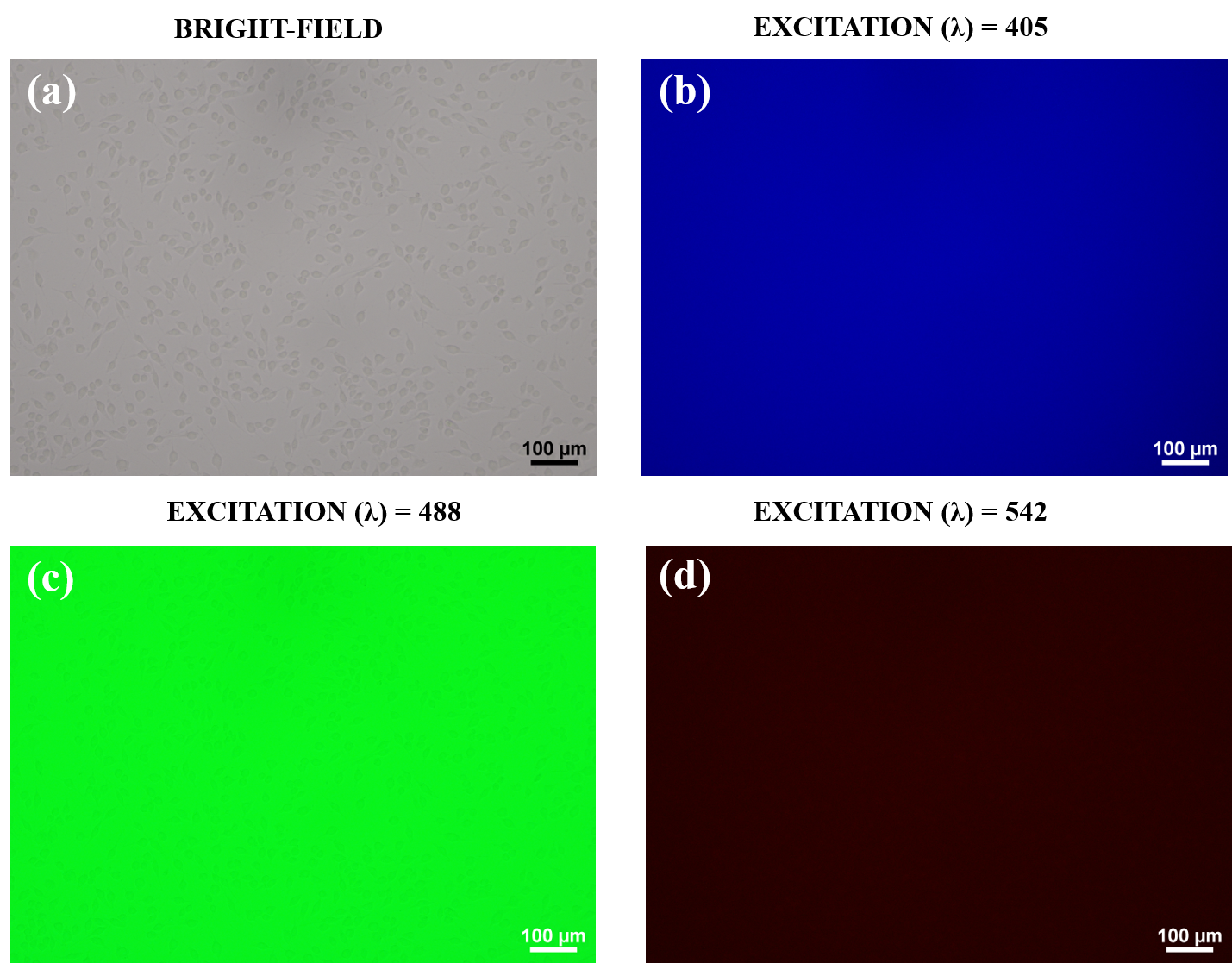

**Fig. S.3** The untreated fibroblast cells (without PPCCDs labelling) excited by laser of wavelength 405 nm, 488 nm, and 542 nm, respectively. Scale bar: 100 µm.

**Quantum yield-** The quantum yield (Q) is ratio of photon absorb to photon emitted which help to determine the comparative yield of desired sample (CDs) with respect to standard sample. The quinine sulfate used as standard sample in the experiment. The quantum yield of PPCCDs and CCDs were calculated using the following equation.

**Quantum Yield (Q) = Q_R_**
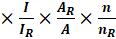

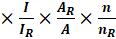

Where, A and A_R_ belongs to absorbance, I and I_R_ is integrated emission intensity, n and n_R_ is refractive index of solvent used for both CDs and reference, and Q_R_ refers the quantum yield of standard quinine sulfate. The calculation of quantum yield of CCDs and PPCCDs were obtain from the slope of **Fig S4** and are shown in **Table S1**.

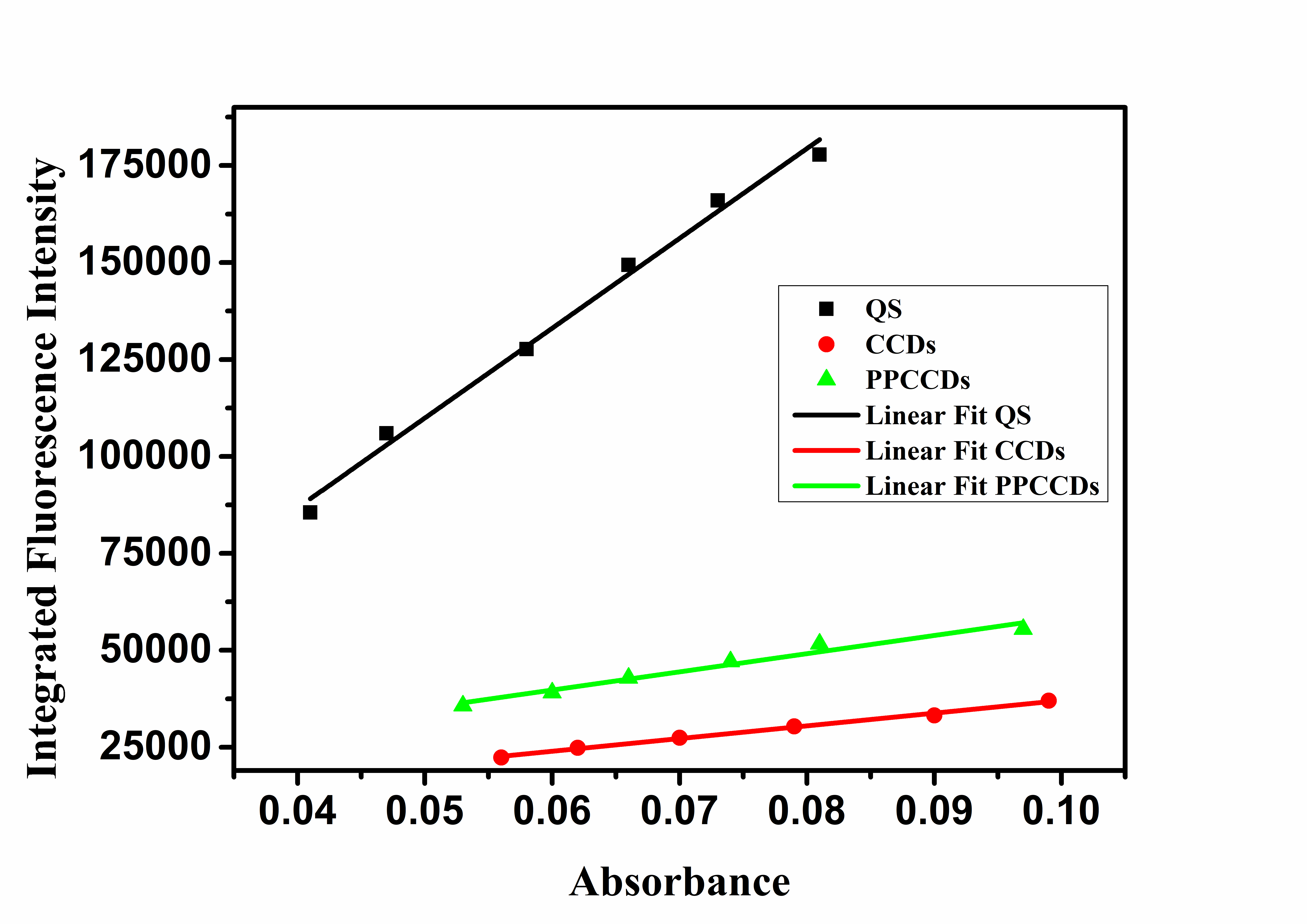

**Fig. S4** Quantum yield plot of PPCCDs and CCDs considering quinine sulfate as standard.

**Table S.1** Calculation of Quantum yield of CCDs and PPCCDs considering quinine sulfate as standard.

| **S. No.** | **Sample** | **Slope** | **Refractive index solvent (ɳ)** | **Quantum Yield (%)** |
| --- | --- | --- | --- | --- |
| **1** | Quinine sulfate | 2315780 | 1.33 | 54 |
| **2** | CCDs | 327486 | 1.33 | 7.6 |
| **3** | PPCCDs | 468711 | 1.33 | 10.9 |

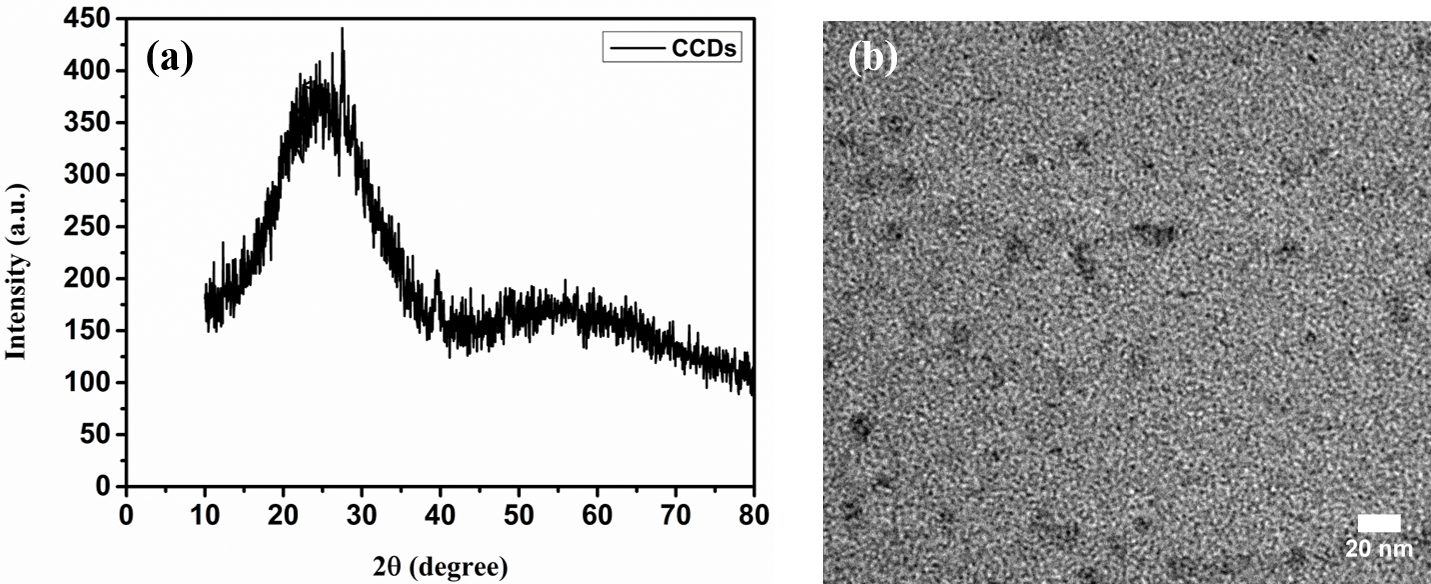

**Fig. S.5** (a) XRD spectra (b) TEM image of clove derived carbon dots (CCDs).

**Table S.2** Comparison of antioxidant efficacy of various carbon dots from derived from different carbon precursors.

| **Carbon precursor** | **DPPH Antioxidant Activity (%)** | **Superoxide Radical Inhibition (%)** | **Ref.** |
| --- | --- | --- | --- |
| Pomelo and ammonium persulfate | 56 | 88 | [1] |
| Pomelo juice and sulfamic acid | 62 | 82 | [2] |
| Proanthocyanidin and ethylenediamine | 30 | - | [3] |
| Polyethyleneimine and cysteine | 74.8 | 68 | [4] |
| Selenocysteine | 70 | - | [5] |
| Black soya beans | 62.8 | 81.3 | [6] |
| **PPCCDs** | **94** | **89** | **This work** |

**Table S.3** Comparison of rate constant performances of various nano-catalyst over degradation of Rhodamine-B

| **Nanomaterials** | **K_app_ (10^-3^ min^-1^)** | **K (s^-1^ g)** | **Ref.** |
| --- | --- | --- | --- |
| Au/CeO_2_-TiO_2_ nano-hybrid | 223.9 | 0.196 | [7] |
| Au nanocrystal  Ag nanocrystal | 0.033  0.007 | - | [8] |
| ZnCO_2_O_4_ nano-dice | 70.4 | - | [9] |
| CDs | 34 | - | [10] |
| N-CQDs | 22.4 | - | [11] |
| **PPCCDs** | **88.7** | **7.39** | **This work** |

**Table S.4** Comparison of various surface passivated carbon dots with their quantum yield and multifunctional aspect.

| **Carbon precursors with passivating agent** | **Methods** | **Quantum yield (%)** | **Applications** | **Ref.** |
| --- | --- | --- | --- | --- |
| Curcumin, PEI | Hydrothermal | 2.28 | Imaging, sensing | [12] |
| Glycerol, (TTDDA) | Microwave | 12.02 | Bioimaging | [13] |
| Chitosan, PEG | microwave | 5.06 |  | [14] |
| Galactose, Polydopamine | microwave | 4.6 | In-vivo imaging | [15] |
| **Clove, PVP** | **Hydrothermal** | **10.9** | **Bioimaging, Antioxidant, catalyst** | **This work** |
